# Quantitative Cell Proteomic Atlas: Pathway-scale targeted mass spectrometry for high-resolution functional profiling of cell signaling

**DOI:** 10.1101/2022.04.14.488331

**Authors:** Paolo Cifani, Alex Kentsis

## Abstract

In spite of extensive studies of cellular signaling, many fundamental processes such as pathway integration, cross-talk and feedback remain poorly understood. To enable integrated and quantitative measurements of cellular biochemical activities, we have developed the Quantitative Cell Proteomics Atlas (QCPA). QCPA consists of panels of targeted mass spectrometry assays to determine the abundance and stoichiometry of regulatory post-translational modifications of sentinel proteins from most known physiologic and pathogenic signaling pathways in human cells. QCPA currently profiles 1,913 peptides from 469 effectors of cell surface signaling, apoptosis, stress response, gene expression, quiescence, and proliferation. For each protein, QCPA includes triplets of isotopically labeled peptides covering known post-translational regulatory sites to determine their stoichiometries and unmodified protein regions to measure total protein abundance. The QCPA framework incorporates analytes to control for technical variability of sample preparation and mass spectrometric analysis, including TrypQuant, a synthetic substrate for accurate quantification of proteolysis efficiency for proteins containing chemically modified residues. The ability to precisely and accurately quantify most known signaling pathways should enable improved chemoproteomic approaches for the comprehensive analysis of cell signaling and clinical proteomics of diagnostic specimens. QCPA is openly available at https://qcpa.mskcc.org.

## Introduction

Targeted mass spectrometry has been established as a highly sensitive, robust, and precise analytical tool for protein and peptide quantification[1–4]. Unlike genomic and transcriptomic analyses, quantitative measurements of peptides and proteins can be used to probe directly the functional differences in signaling pathways across cell states and types, providing a powerful tool to reveal the effects of functional perturbations that occur physiologically or in disease. The availability of panels of targeted peptide assays, such as those developed in the CPTAC Assay Portal[5–8], has been critical for establishing these technologies and their use. Peptides included in these panels are selected for their sequence specificity and for having chemical and physical properties suitable for mass spectrometric detection, thus permitting highly sensitive and specific measurements of complex biological samples[9].

While cellular concentration is an important determinant of protein activity, post-translational chemical modification is pervasive and equally important mode of biochemical regulation of protein activity. For example, biological functions of protein phosphorylation have been documented extensively, including causal linkage of aberrant activity of protein kinases and phosphatases to cancer onset and progression[10–12]. Systematic quantification of specifically phosphorylated regulatory protein domains has been used to determine cellular functional states[13] and to identify molecular mechanisms linked to disease progression, and response and resistance to therapy [13–18]. To aid the detection of phosphorylated proteins and peptides, several immune- and metal affinity enrichment techniques have been recently developed [19–22].

However, affinity enrichment is not absolutely required for the detection of chemically modified peptides. For example, we and others have used high-resolution multidimensional liquid chromatography to measure modified and unmodified peptides without enrichment per se [23,24]. This can be combined with targeted precursor ion quantification to measure molecules with fragmentation properties that are not robust enough for reaction monitoring techniques, as frequently observed for phosphorylated peptides[25]. Combined with the recently developed methods for multiplexing, such as isobaric labeling and triggered scanning [26–28], these approaches can be used for comprehensive, integrated, and quantitative analyses of cell signaling.

To enable pathway-scale targeted mass spectrometry for high-resolution functional profiling of cell signaling, we have now developed the Quantitative Cell Proteomic Atlas (QCPA) to determine abundance and stoichiometry of regulatory post-translational modifications of proteins covering most known physiologic and pathogenic signaling pathways in human cells. These processes include cell surface signaling, regulation of cell proliferation and death, homeostatic and stress responses, regulation of gene expression, among others. We incorporated them into an openly accessible and interactive database, designed for modular future growth and integration with other public data (https://qcpa.mskcc.org/). As part of this framework, we also developed *TrypQuant*, a synthetic spike-in reagent for rapid and precise quantification of trypsin proteolysis efficiency in order to control for technical variability of sample preparation and mass spectrometric analysis. We anticipate that QCPA and related approaches to precisely and accurately quantify most known signaling pathways should enable improved chemoproteomic approaches for the integrative analysis of cell signaling and implementation of functional biomarkers for clinical proteomics.

## Experimental section

### Design of QCPA assays

All QCPA peptide assays were designed and refined individually. Proteins and peptides were chosen based on published biological activities (see Results and Discussion section). Assays were designed using the canonical protein isoforms as per UniProt database[29]. All assays were designed assuming tandem protein proteolysis using LysC and trypsin.

For peptides containing two known post-translational modification (PTM) sites (site “A” and “B”, respectively), all three modification configurations were included (i.e. modification of site A, of site B, and of sites A+B). Peptides containing more than two PTM sites were omitted. Peptides were also omitted if they included less than seven or more than thirty residues, and if mapping to the protein termini. Peptides flanked by ragged tryptic sites (i.e. RR, RK, KR, KK) were omitted unless covering main regulatory sites, or if alternative known functional PTM sites were not available for that protein. In this case, assays were designed assuming tandem proteolysis with LysC and trypsin, and thus favoring the lysine residue as primary cleavage site. Specific proteolysis by cellular signaling proteases (e.g. caspases) was conceptually equated to a functional PTM. Consequently, semitryptic peptides resulting from sequence-specific proteolysis were included if at least the non-modified peptide fulfilled the criteria above. QCPA assays were restricted to proteotypic peptides, as determined by comparing the sequences with canonical human proteomes annotated both in the UniProt and in the NCBI RefSeq databases[29,30]. The uniqueness criterion was relaxed for degenerate peptides covering conserved regulatory sites in functionally related isoforms (e.g. main regulatory site of RSK1, RSK2, and RSK3 kinases, see Figure 2).

Peptides for the quantification of unmodified peptides and proteins were chosen to be proteotypic, to contain 10-15 residues with no missed tryptic cleavages, and to be frequently observed by mass spectrometric analysis using data reported in PeptideAtlas [31]. Furthermore, assays for protein quantification were required to be generated by non-ragged tryptic sites, and to contain no RP or KP motifs. To reduce the effects of experimental chemical modifications, peptides containing methionine, glutamine, or asparagine residues, or more than one histidine residue, were omitted. Peptides with repetitive sequences and peptides containing more than one proline residues were also omitted, as these features are challenging for solid-phase peptide synthesis and thus potentially limit the use of QCPA for applications requiring synthetic reference standards[1,32,33].

### TrypQuant

The TrypQuant standard was designed to include two unmodified tryptic sites (one K and one R), one monomethyl-R residue, one acetyl-K residue, and two unmodified sites (one K and one R) adjacent to a phosphorylated residue. The sequence was designed to have no peptides mapping to the canonical human and mouse proteomes (UniProt and RefSeq databases as of July 2019), and to contain no C, N, F, M, P, and H residues. Each product peptide was designed to contain one Y residue to enable spectrophotometric quantification. Peptides were designed to have m/z in the 380-690 Th range and to be doubly charged. The TrypQuant reagent consist of the following peptides (Figure 1A).

**Figure 1:**
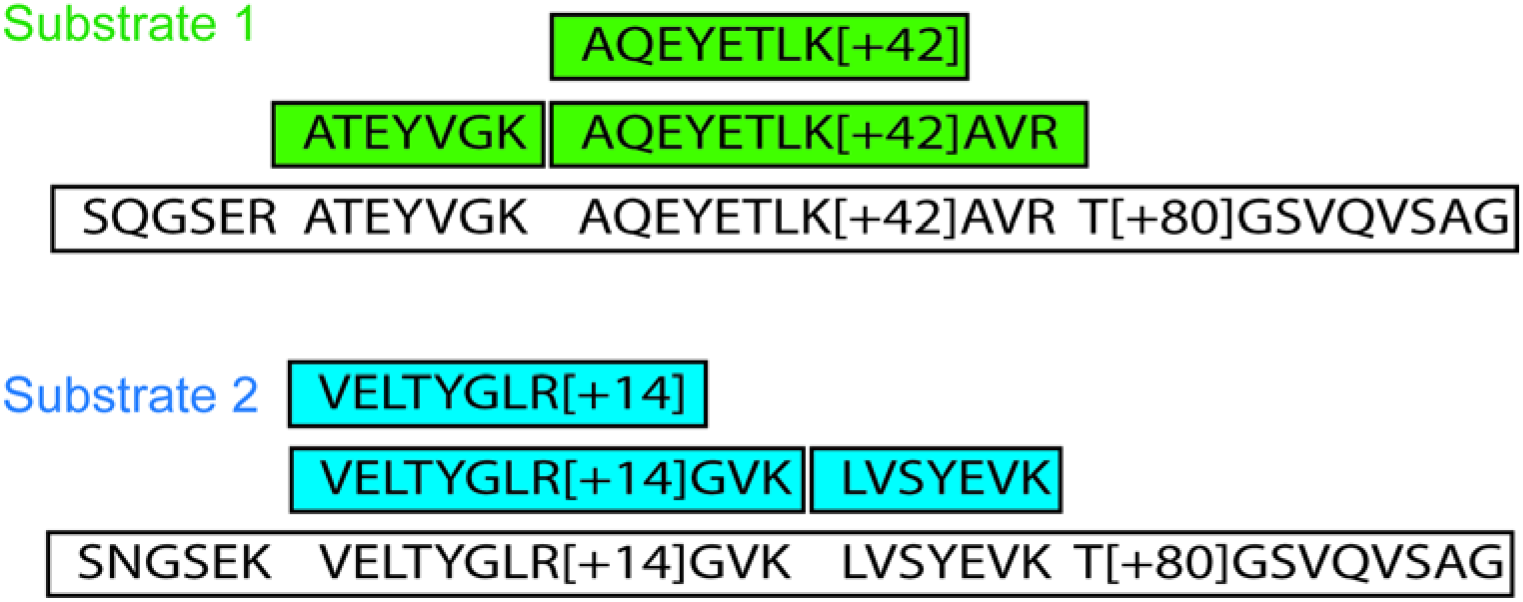
Representation of TrypQuant components. The substrate peptides are rendered in white, with chemically modified sites indicated as “+14” for methylation, “+42” for acetylation, and “+80” for phosphorylation, respectively. The measured product peptides for Substrates 1 and 2 are labeled in green and blue, respectively.

### Synthetic peptides

QCPA peptides were synthesized by JPT Peptide Technologies GmbH (Berlin, Germany) as SpikeMix Peptide Pools (purity>75%), in sets of 96 peptides per pool. ^13^C_6_^15^N_2_-lysine and ^13^C_6_^15^N_4_-arginine residues were incorporated as C-terminal amino acids. All C residues were carbamidomethylated. Sets of lyophilized peptides were solubilized at 100 pmol/μl in 30% aqueous acetonitrile by sonication for 5 minutes, pooled, and stored at −80C. Immediately before analysis, the synthetic peptide library was thawed on ice and diluted in MV4-11 tryptic cells proteome (prepared as described in [23]) to obtain 250 fmol/μl synthetic peptides and 2.5 μg/μl cell line proteome in 0.1% formate in water.

TrypQuant peptides were synthesized by New England Peptides (Gardner, MA, USA) as individual peptides (purity >95%). Substrate peptides were labeled with ^13^C_6_^15^N_2_-lysine and ^13^C_6_^15^N_4_-arginine residues, while product peptides contain amino acids with natural isotopic distribution. Peptides were reconstituted in in 20% aqueous acetonitrile by sonication for 5 minutes, quantified based on their absorbance at λ=280 nm using Nanodrop 2000 spectrophotometer (Thermo Fisher), and pooled at equimolar amounts. After overnight digestion with Sequencing Grade Modified Porcine Trypsin (Promega, Madison, WI, USA), peptides were quantified by label-free LC-MS, and observed deviations from equimolarity were corrected based on these empirical measurements (Figure 1B).

### Chromatography, mass spectrometry and MSMS analysis

To verify the ionization properties of QCPA peptides, 500 fmol QCPA library diluted in 5 μg purified MV4-11 tryptic proteome was resolved by nano-scale multidimensional chromatography, ionized by electrospray ionization, and analyzed by targeted MSMS. A detailed description of the chromatographic system and the electrospray ion source is provided in [23]. In short, liquid chromatography experiments were performed using an Ekspert NanoLC 425 chromatograph (Eksigent, Redwood City, CA, USA), equipped with an autosampler module, two 10-port and one 6-port rotary valves, and one isocratic and two binary pumps. Peptides were loaded on a 100μm ID x 150 mm capillary strong-cation exchange column packed with Polysulfoethyl A 5 μm silica particles (PolyLC, Columbia, MD, USA). Peptides were eluted by a step gradient of aqueous ammonium formate pH3 (15, 30, 62.5, 125, 250, 500, 1000 mM, plus a final aliquot of 1M ammonium formate pH 9) into a reversed phase trap column (100μm ID x 50 mm, packed with Poros R2 10 μm C18 particles (Life Technologies, Norwalk, CT, USA) using a vented trap-elute configuration. Eluates from each SCX fraction were resolved on a analytical reversed phase capillary column (75 μm x 500 mm, packed with Reprosil 1.9 μm silica C18 particles (Dr. Meisch, Ammerbauch-Entrigen, Germany), kept at constant 60°C. Chromatographic separation was performed using a 150 min 3-35% acetonitrile gradient in water (0.1% formate) at a flow rate of 250 nl/min. The reversed phase column was directly connected to 3 μm ID electrospray emitter[34] using a metal union that also served as the electrospray electrode. Electrospray ionization was achieved using variable voltage, programmed from 1750 to 1450 V with 50 V steps over the gradient elution using a DPV-565 PicoView ion source (New Objective, Woburn, MA, USA).

Spectra were acquired using Orbitrap Fusion mass spectrometer (Thermo Scientific, San Jose, CA, USA). The mass spectrometer was set to perform a precursor scan (350-2000 Th) every 3 s at 240,000 resolution. Ions were selected for fragmentation using the “Trigger ion” routine (10 ppm tolerance), with m/z inclusion list compiled based on the expected mass and charge. When the number of trigger peptides was insufficient to fill the time between precursor scans, conventional data-dependent acquisition was programmed. Fragmentation spectra were collected in the linear trap at “normal” scan rate (unit resolution) after HCD fragmentation at 28, 32, and 38% stepped normalized collision energy, with AGC set to 10,000 ions and max injection time set to 100 ms. Peptides not detected using this method were included in a targeted inclusion list which included the expected m/z for z ranging from 1 to 5 (only if the precursor m/z was included in the 350-2000 Th range), and fragmentation of these peptides was programmed using the targeted MS2 (i.e. PRM) routine.

Fragmentation spectra were searched against the UniProt human database, supplemented with the sequences of mutant proteins included in QCPA (e.g., KRAS^G12D^) using in MaxQuant and PEAKS [35,36], with mass tolerance set to 5 ppm and FDR<0.05. TrypQuant peptides were quantified by integrating the elution profiles of the precursor ions using Skyline[37]. All methods conform to the National Institute’s of Health Rigor and Reproducibility Principles and Guidelines.

## Results

The Quantitative Cell Proteomic Atlas (QCPA) is designed to enable systematic and integrated measurements of protein abundance and stoichiometry of post-translational modifications by targeted mass spectrometry [37]. Based on this rationale, QCPA organizes peptide assays by signaling pathways and their biochemical surrogates. While the current release of QCPA includes most known signaling pathways, QCPA also includes biochemical processes reported to contribute to disease development and modulate therapy response, i.e. pathways mediating response to cellular stress and drug metabolism (Figure 1A & S1)[38].

For each of these processes, relevant biochemical pathways, proteins and biochemical surrogates were identified and prioritized based on the review of published literature. For selected pathways, such as the RAS-RAF-MEK-ERK kinase signaling pathway, additional activity modulators, including protein phosphatases and co-activators, were included to enable high-resolution molecular profiling (Figure 2). The modular organization of QCPA is designed to be easily expanded to include additional biochemical processes and pathways in the future. With this goal in mind, QCPA implements a collaborative approach to facilitate expert curation and review to suggest the inclusion of future pathways, using an online user-controlled database, as implemented in https://qcpa.mskcc.org/.

**Figure 2:**
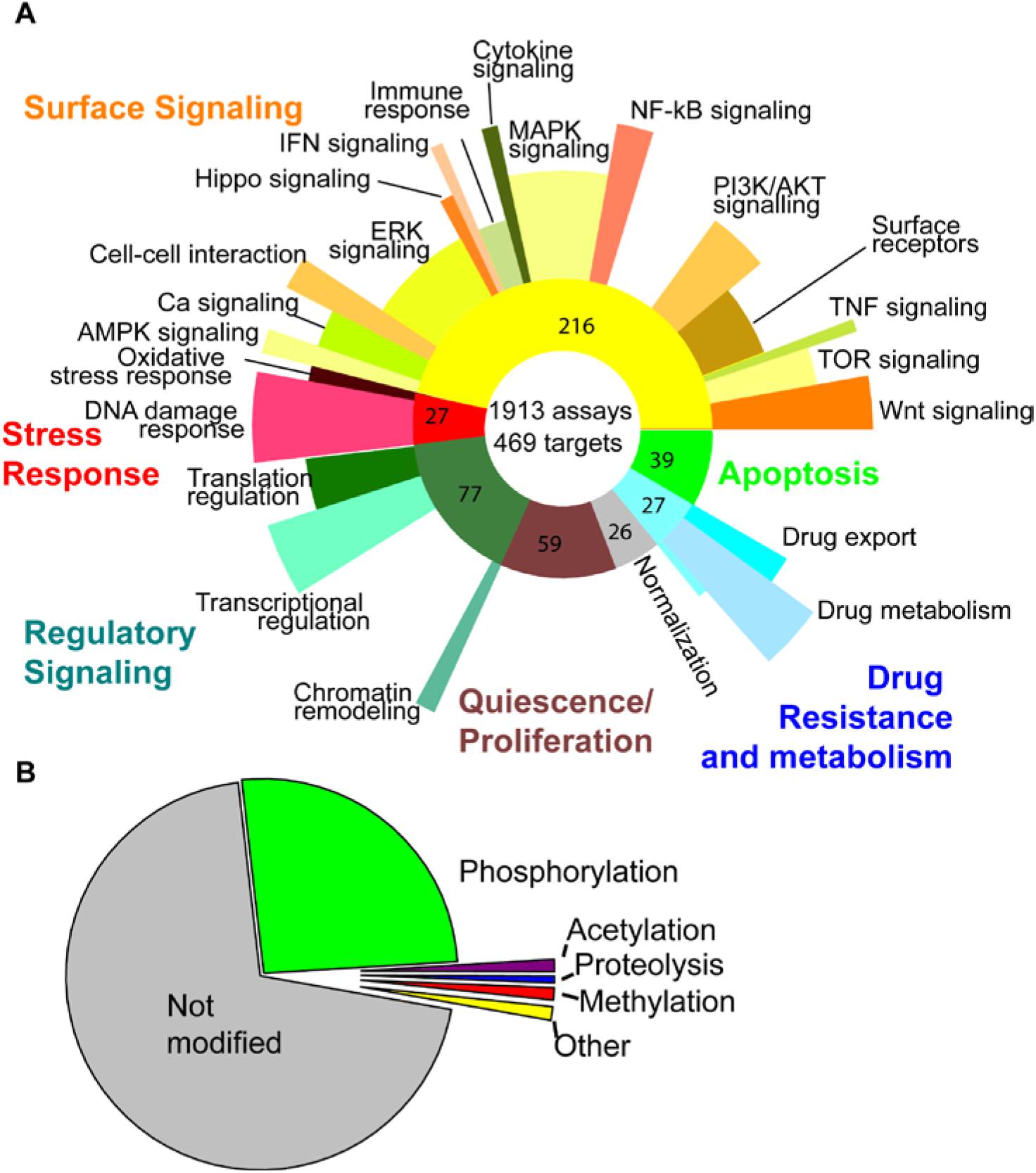
A. Overview of QCPA biochemical processes and pathways. The number of peptide assay for each category is reported in the correspondent portion of the graph. B. Chemical modification status of peptides in current QCPA version.

QCPA is a repository of peptide assays designed for quantitative targeted mass spectrometry. QCPA assays can be implemented on diverse mass spectrometers, including ion trap and time-of-flight instruments, and using various quantitative precursor and fragment ion-based approaches[22]. Each peptide was individually designed to be compatible with electrospray ionization and avoid known artifacts in sample or ion stability such as inefficient proteolysis and ion fragmentation. We used the sentinel concept to define key proteins for specific signaling pathways, based on the review of published literature[15]. For proteins with regulatory PTMs, regulatory sites were defined based on published experimental evidence, including those established by immunoassay and proteomics methods [39]. Sites without established functions, based on literature review curated from PubMed as of 2018, were not considered in the current implementation of QCPA.

Because kinase and phosphorylation signaling are among the most studied mechanisms[40–42], approximately 25% of QCPA assays focus on serine, threonine, and tyrosine phosphorylated peptides (Figure 2B). Additional regulatory PTMs include lysine acetylation and methylation, as well as site-specific proteolysis. QCPA is organized modularly to permit additional and novel PTMs to be readily incorporated in the future. Cellular protein concentration is a known determinant of biochemical activities and cellular states[43]. For all proteins whose PTM regulation is profiled by QCPA assays, QCPA also includes two unique proteotypic peptides per protein without known modifications designed to measure differential protein abundance (Figure 3). This triplet framework allows for direct measurements of protein and peptide abundance as well as PTM stoichiometry[44].

**Figure 3:**
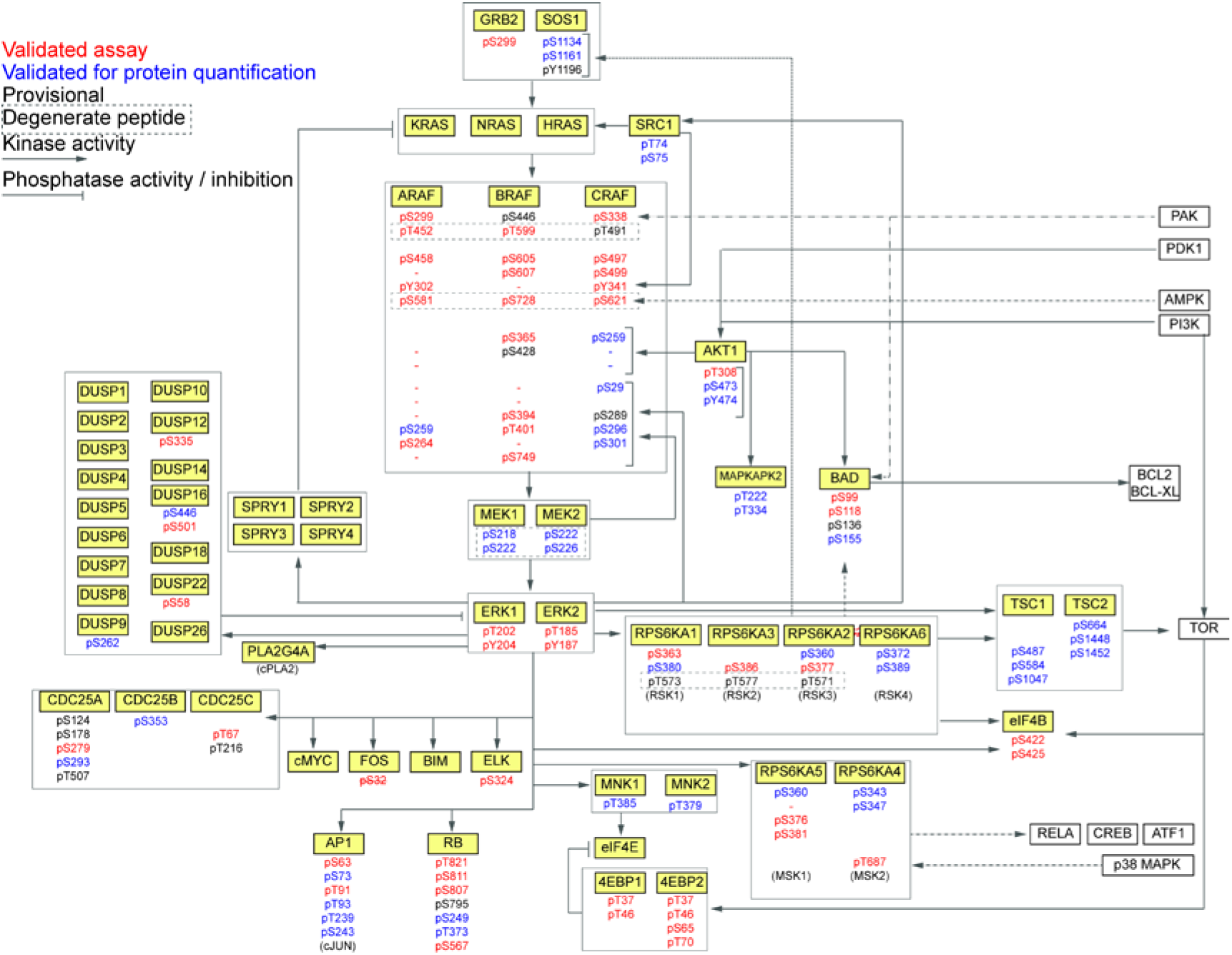
Schematic of QCPA assays to profile the RAS-RAF-MEK-ERK signaling pathway. Validated and provisional assay peptides are marked in red and blue, respectively. Arrows and lines indicate publicly reported activating and inhibitory interactions, respectively.

In order to confirm the detectability of designed QCPA peptides, we synthesized libraries of isotopically-labeled peptides, diluted them in whole-cell tryptic proteome isolated from human cells, and analyzed the resultant mixtures containing 500 fmol of QCPA peptides by two-dimensional chromatography and targeted high-resolution Orbitrap mass spectrometry. This confirmed the detection of 1,577 QCPA peptides, which we term validated, with 336 additional provisional QCPA peptides requiring additional optimization in the future. On average, validated QCPA peptides covered 84% of sentinel proteins per pathway (range from 72 to 100%; Figure 1), including 71% of phosphorylated peptide assays (range 42 to 100%). For example, validated QCPA peptides provide near complete measurements of all known PTM and protein regulation of the RAS-RAF-MEK-ERK signaling pathway (Figure 3).

To enable comparative studies of different tissues and cell types, QCPA includes 26 specific peptides for specimen and tissue normalization [45]. They were designed to measure the abundance of 23 distinct proteins from different subcellular compartments, and thus can also serve as measurements of efficiency of specimen and proteome extraction. For example, this includes nuclear NCL, H1B, and PCNA proteins, cytoskeletal ACTB and VIM, and mitochondrial ATP5B, ER CALR, and Golgi TGOLN2, among others.

QCPA is a peptide-centric atlas, dependent on the complete proteolysis of target proteins for accurate mass spectrometric quantification. Because this process is sample and condition dependent, and additionally affected by post-translational modifications adjacent to protease substrate, we designed a synthetic reagent, termed *TrypQuant*, to be added and proteolyzed with the sample proteome. This substrate contains both lysine and arginine residues positioned next to phosphorylated, methylated and acetylated amino acids (Figures 4 and 5). *TrypQuant* is composed of two equimolar and isotopically labeled peptide pairs: two concatenamer substrates, and their tryptic product peptides. Proteolysis of the substrates produces peptides that can be quantified relative to their reference isotopologues based on the intensity of their respective precursor ions. To confirm this experimentally, we generated an equimolar mixture of concatenamers and product peptides based on their UV absorbance (see Methods section). Proteolysis was performed to completeness, and the amount of each TrypQuant component was recalibrated based on observed MS ion currents. Recalibrated TrypQuant was then co-digested with a cell line protocol under conditions typical for proteomic experiments. As a result, the extent of tryptic proteolysis can be precisely quantified for each sample and used as a correction factor, if appropriate. For example, this analysis showed essentially no apparent proteolysis of acetylated lysine residues, but residual activity on monomethylated arginine substrates (Figure 5C).

**Figure 4:**
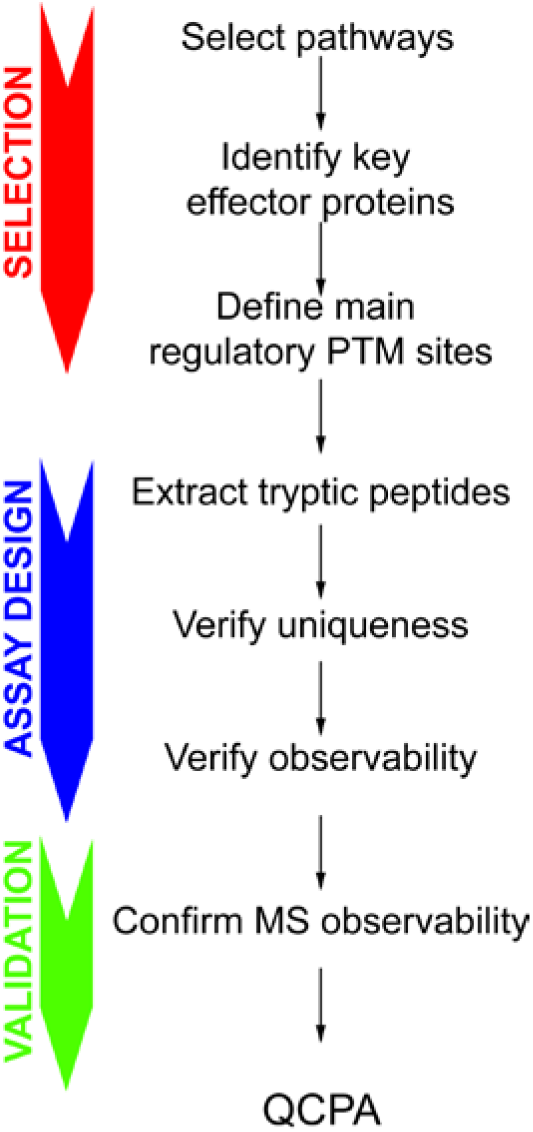
Schematic of the QCPA assay design and selection.

**Figure 5:**
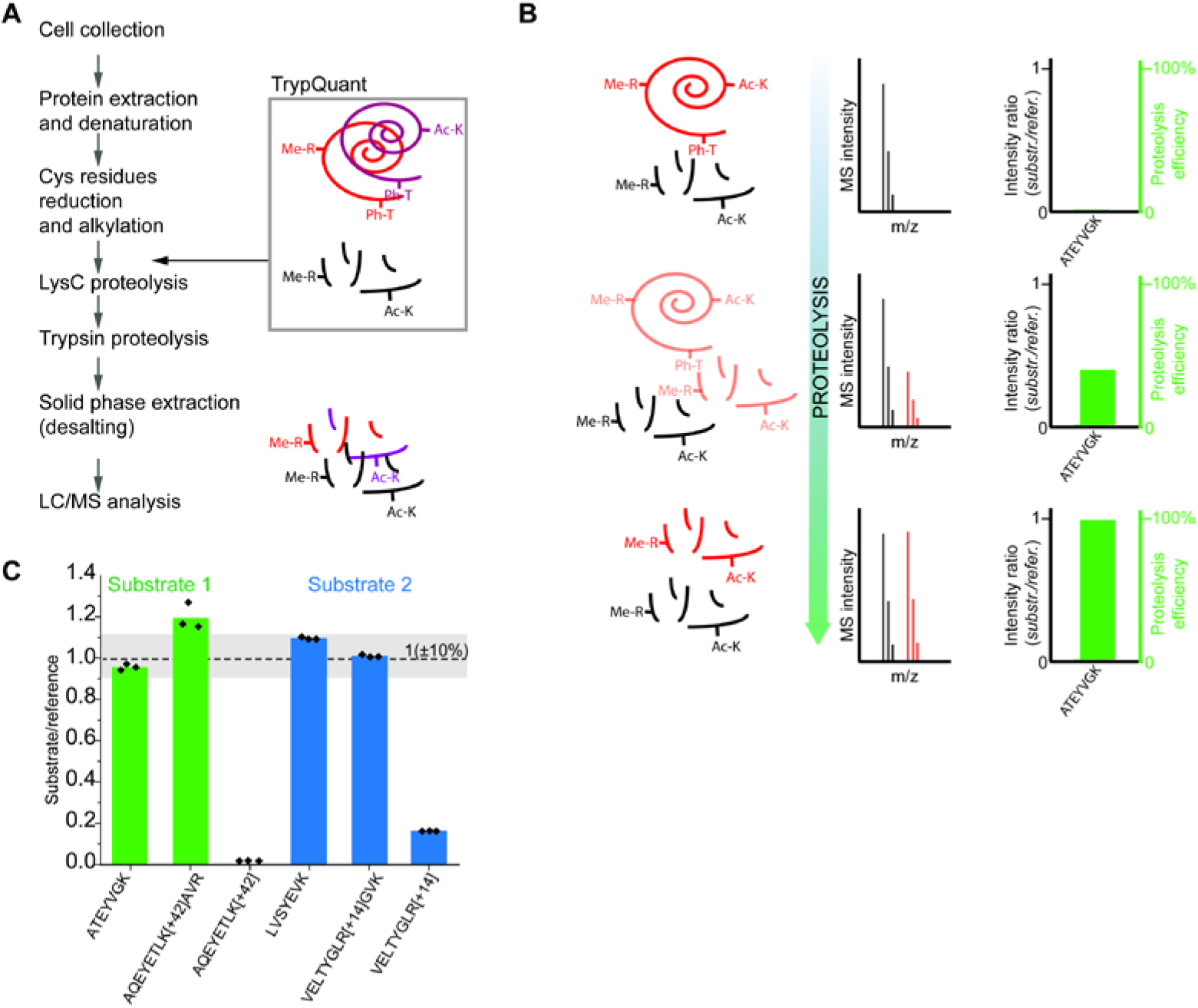
Overview of the TrypQuant design and use. A. Synthetic TrypQuant peptides are added to the sample before processing and proteolysis, and the substrate concatenamers are digested together with the sample proteome, generating isotopically labeled product peptides. B. The relative ion intensity of product peptides and reference directly quantify the degree of proteolysis for different substrates. C. Results of the empirical recalibration of TrypQuant components.

The QCPA database has a peptide-based structure, where every peptide is annotated with a unique QCPA identifier, UniProt ID and name of the corresponding protein, official and alternative gene names, amino acid sequence, peptide modification site as per canonical isoform in UniProt, and functional category. For peptides that are also currently included in the CPTAC Assay Portal (145 peptides corresponding to 9% of validated QCPA assays), QCPA database also lists the corresponding CPTAC identifiers (https://proteomics.cancer.gov/assay-portal). QCPA peptide assays are also annotated with relevant publications describing the protein function. QCPA is openly accessible via an interactive web interface, including search and browse functions to identify available assays for pathways or proteins of interest (https://qcpa.mskcc.org).

## Conclusions

Here we introduce the Quantitative Cell Proteomic Atlas for pathways-scale targeted mass spectrometry for high-resolution functional profiling of cell signaling. QCPA is organized in panels of targeted mass spectrometry assays for the quantification of differential biochemical regulation across cell states. This approach is explicitly designed to measure differential regulation both due to changes in protein abundance and stoichiometry of post-translational modifications of functionally established regulatory sites. This should pave the way to integrative approaches for systems biology and new diagnostic tools for clinical use [46].

The rationale for the development of QCPA assays is eminently biological. As such, QCPA assays can be used with both high-resolution ion trap and time-of-flight mass spectrometers, including diverse quantitative targeted methods such as SureQuant [19], AIM [23] and others. In addition, the use of QCPA libraries of isotopically labeled peptides permits isotopically-triggered scanning and other automated mass spectrometry acquisition methods [17,28,47,48] to increase assay multiplexing and throughput [4,49]. While specific methods and applications will require explicit optimization of the analytical performance [7,50], the breadth and depth of QCPA targets should enable increasingly comprehensive and integrated measurements of cell signaling.

The availability of targeted peptide assays specifically designed to quantify functional biochemical states should also permit measurements of signaling in rare cell populations, including single cells [51][52]. Finally, QCPA paves the way to leveraging targeted profiling of biochemical activation as an objective parameter for clinical diagnosis and personalized therapy selection.

## Supporting information

Graphical overview of the proteins and pathways included in QCPA.

List of QCPA assays, as of version 1.2.

List of empirically observable QCPA peptides.

## Acknowledgements

AK is a Scholar of the Leukemia & Lymphoma Society and acknowledges support from NCI R01 CA204396, R21 CA235285, and P30 CA008748. The authors thank Slav Maron for developing the QCPA database and online interface.

## Supporting Information

**Supplementary Figure 1:** Graphical overview of the proteins and pathways included in QCPA.

**Supplementary Table 1:** List of QCPA assays, as of version 1.2.

**Supplementary Table 2:** List of empirically observable QCPA peptides.

