## Supplementary figures and images for "Quantitative Cell Proteomic Atlas: Pathway-scale targeted mass spectrometry for high-resolution functional profiling of cell signaling"

### Graphical overview of the proteins and pathways included in QCPA.

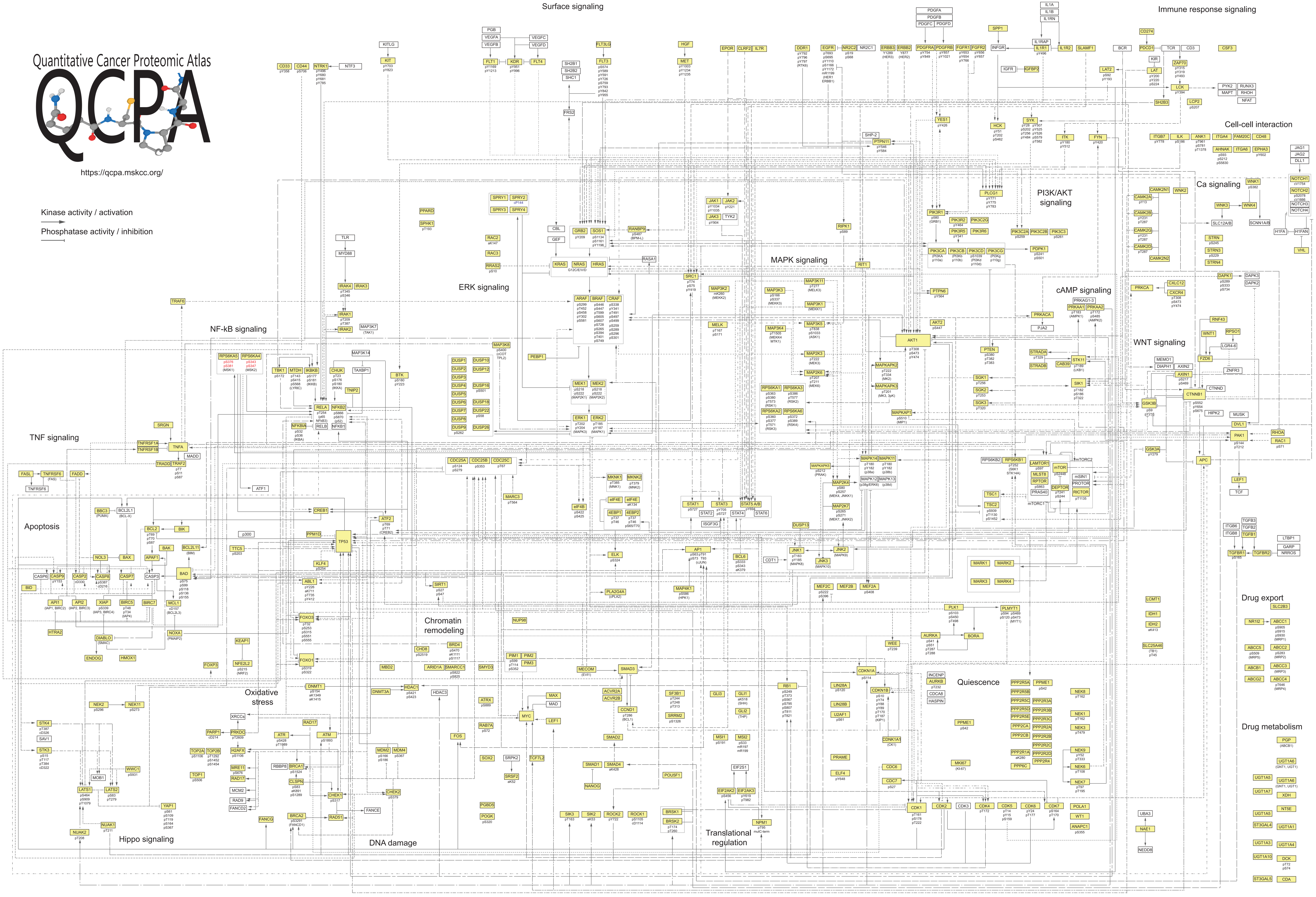
